# A Novel Proteoglycan-4 Isoform Drives Skeletal Regeneration

**DOI:** 10.1101/2025.09.15.676417

**Authors:** Mohamed Tantawy, Deepak K. Khajuria, Christopher C Norbury, Fadia Kamal, Reyad A. Elbarbary

**Affiliations:** Department of Orthopaedics and Rehabilitation, The Pennsylvania State University College of Medicine, Hershey, Pennsylvania 17033, USA; Center for Orthopaedic Research and Translational Science (CORTS), The Pennsylvania State University College of Medicine, Hershey, Pennsylvania 17033, USA; Department of Cell and Biological Systems, The Pennsylvania State University College of Medicine, Hershey, Pennsylvania 17033, USA; Center for Cannabis and Natural Product Pharmaceutics (CCNPP), The Pennsylvania State University College of Medicine, Hershey, Pennsylvania 17033, USA; Department of Molecular and Precision Medicine, The Pennsylvania State University College of Medicine, Hershey, Pennsylvania 17033, USA; Center for RNA Molecular Biology, Pennsylvania State University, University Park, Pennsylvania 16802, USA

**Keywords:** Fracture healing, bone regeneration, proteoglycan, alternative splicing, stem cell

## Abstract

Proteoglycan 4 (PRG4) is an extracellular matrix protein best known for its lubricating role in articular cartilage. Using a murine model of tibial fracture, we performed RNA-seq across multiple post-fracture timepoints and observed high *Prg4* expression during the first week of healing, coinciding with the initial inflammatory phase. Single-cell RNA-seq of the fracture callus localized *Prg4* expression to a stem cell population with key roles in repair. Analysis of alternative splicing in callus RNA identified a unique *Prg4* isoform lacking three coding exons (hereafter termed *Prg4-S*). In vivo knockdown of *Prg4-S* using locally delivered siRNA produced multiple defects that culminated in impaired bone formation. Together, these findings uncover an unanticipated osteogenic role for a unique *Prg4* splicing isoform and highlight its potential for translational applications.

## Introduction

The repair of most of the cortical bone fractures proceeds through three main phases: inflammation, repair, and remodeling^1,2^. During the inflammatory phase, immune cells invade the fracture gap and secret cytokines and growth factors that recruit other immune cells, as well as stem cells and bone progenitors, to set the stage for the repair phase^1,2^. During the early repair phase, the fracture gap is filled with a fibrocartilaginous (soft) callus composed mainly of proliferating chondrocytes^1,2^. These chondrocytes later undergo hypertrophy and mineralization, which hardens the fracture gap and promotes formation of new blood vessels across the fracture line^1,2^. The mineralized soft callus is then replaced by bone, forming a bony callus at the end of the repair phase. Finally, the healing process concludes with remodeling of the newly formed woven bone to re-establish the characteristic laminar structure of the cortical bone^1,2^.

The extracellular matrix (ECM) is an integral component of bone tissue. Proteoglycans are a major constituent of the bone ECM and play important roles in maintaining tissue integrity^3^. Proteoglycans consist of a core protein covalently attached to multiple chains of linear polysaccharides, glycosaminoglycans (GAGs)^4^. GAGs add structural complexity to proteoglycans and facilitate interactions with other ECM components. Owing to their hydrophilic nature, GAGs enable proteoglycans to attract and retain large amounts of water, thereby maintaining tissue hydration^4^. Members of the proteoglycan family differ in their core proteins as well as in the type and size of their attached GAGs. Proteoglycan 4 (*Prg4*; also known as lubricin), a member of this family, is conserved across the animal kingdom^5^. It is highly expressed in vertebrate synovial fluid and in the superficial zone of articular cartilage, where it functions both as a structural protein and as a lubricant, reducing shear forces at the cartilage surface^5–8^. *Prg4* exerts chondroprotective effects, and deficient *Prg4* expression in the joint is associated with the onset of osteoarthritis (OA)^9–11^. Consequently, recombinant PRG4 protein has been proposed as a disease-modifying therapy for OA^6,12–17^. Most published functional studies have investigated the biological roles of *Prg4* in cartilage. However, *Prg4* is also expressed in other tissues, where it may play additional roles^5^. Supporting this, loss-of-function mutations in the *PRG4* gene cause the autosomal camptodactyly-arthropathy-coxa vara-pericarditis (CACP) syndrome^18–20^, which is characterized by chronic inflammation and pericarditis in addition to joint degeneration^20^. Recent studies have begun to uncover roles for *Prg4* in skin and ear wound closure that are independent of its lubricating function^21–23^. Importantly, the expression patterns and biological functions of *Prg4* during bone healing remain completely uncharacterized.

Alternative splicing (AS) is a fundamental regulatory process that removes introns from precursor (pre)-mRNAs and joins exons in different combinations, thereby generating diverse mature mRNAs from the same gene^24^. AS is a tightly regulated mechanism that plays critical roles in maintaining normal tissue homeostasis^25^. It is tissue- and context-specific, with some AS events occurring only in response to particular signals such as developmental or pathological cues^26^. Despite its importance, AS remains under-investigated in the context of bone repair, and our knowledge of its role within the healing callus is extremely limited. Notably, cytokines and growth factors are known to influence and crosstalk with AS^2,27–31^, and these factors are secreted at high levels during the inflammatory phase of fracture healing—further emphasizing the importance of studying AS in bone repair. Different splicing isoforms of *Prg4* have been characterized in articular cartilage and found to be differentially expressed between the anterior load-bearing and posterior non-load-bearing regions of the femoral medial condyle^32^. However, the functional basis of this anatomical distribution remains unclear. Importantly, all published studies reporting on the biological roles or therapeutic potential of PRG4 have focused almost exclusively on the longest isoform, leaving the other isoforms completely unstudied^21–23^.

Here, we investigated the expression pattern of *Prg4* during fracture healing at the single-cell level and characterized the isoforms of the *Prg4* transcript that are specifically expressed in the callus microenvironment. We also performed functional studies to knock down specific *Prg4* isoforms locally in the callus and defined the roles of *Prg4* in diverse biological processes central to the healing process.

## Results

### Callus progenitors express high levels of *Prg4* during the inflammatory phase

We recently reported bulk RNA-seq data spanning the full course of fracture healing in a murine tibial mid-diaphyseal osteotomy model stabilized with an intramedullary nail^27,33^ (Fig. 1A). Analysis of *Prg4* expression revealed high levels during the initial inflammatory phase (particularly on days 3–5), followed by a steady decline during the cartilaginous phase (days 7– 14) and near absence at later stages (Fig. 1B). The hematoma at days 3-5 is populated mainly by stem/progenitor and immune cells, none of which are known to express *Prg4* at the levels observed in our bulk RNA-seq data. To localize *Prg4* expression to specific cell types, we performed single-cell RNA sequencing (scRNA-seq) on day 5 (d5) and day 10 (d10) callus tissues. Across both timepoints, 32 distinct clusters were identified (Fig. 1C; Supplementary Fig. 1A, B). Among these, cluster 2 (C2) was the primary source of *Prg4* (Fig. 1D). Based on expression of canonical markers including *Prrx1, Pdgfrα,* and *Ly6a (Sca1)* (Fig. 1E), C2 was classified as skeletal stem cells/osteochondral progenitors^34^. C2 also expressed *Ctsk* and *Postn* (Fig. 1E), genes known to mark periosteal stem cells^35–37^. Hereafter, we refer to these cells as Postn⁺ stem cells (PSCs) to distinguish them from other clusters

**Figure 1.**
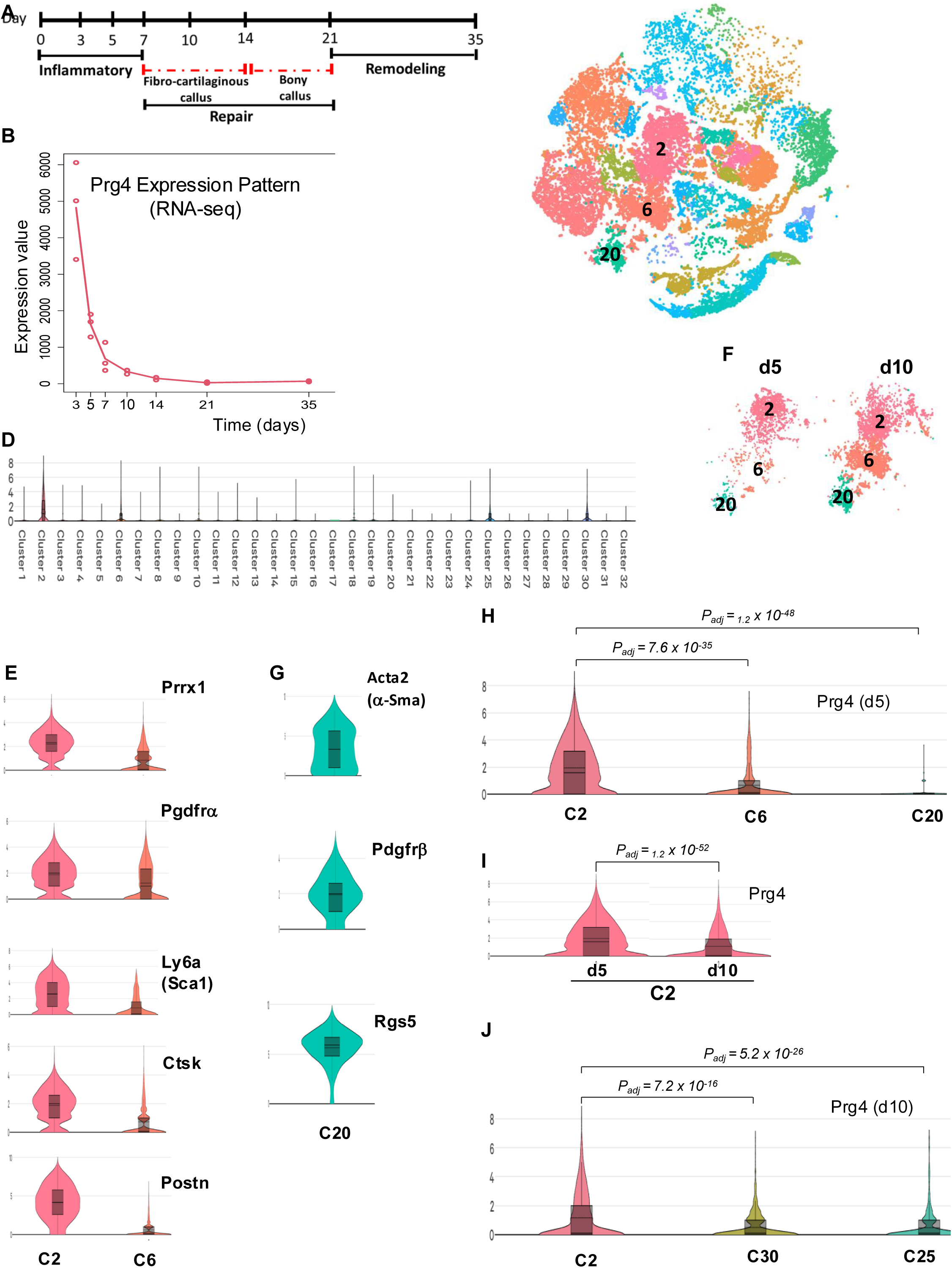
PSCs express *Prg4* during fracture healing. (**A**) Timeline of the healing course, showing the healing phases and harvest time points used for bulk RNA-seq. (**B**) RNA-seq analysis showing Prg4 expression at the indicated post-fracture time points (N = 3). (**C**) Cell clusters identified using scRNA-seq on days 5 and 10 combined. Three mice were used per day, and 22,854 cells were sequenced (median 2,207 genes per cell) (**Supplementary Fig. 1A, B** show each day separately). (**D**) scRNA-seq violin plots showing *Prg4* expression in the 32 cell clusters shown in (**C**). (**E**) scRNA-seq violin plots showing the expression of the specified genes in the indicated clusters. (**F**) Stem cell/progenitor and stem-cell-like clusters identified on days 5 and 10. (**G**) Same as in (**E**). (**H–J**) scRNA-seq violin plots showing *Prg4* expression in the specified clusters on day 5 (**H**), days 5 and 10 (**I**), or day 10 (**J**). Adjusted p-values (Padj) were calculated using the Wilcoxon rank-sum test followed by Benjamini–Hochberg correction.

In addition to C2, clusters C6 and C20 also contained progenitor or mesenchymal stem cell (MSC)-like populations (Fig. 1C, F). C20 was identified as pericytes^38^ (Fig. 1G). Cells in C6 were less well defined: they expressed little to no *Pdgfrα, Sca1, Ctsk,* or *Postn* (Fig. 1E), but variably expressed markers associated with mesenchymal progenitors, MSC-like cells, and activated fibroblasts (Supplementary Fig. 1C). This suggests that C6 represents a heterogeneous pool of mesenchymal-like progenitors distinct from PSCs in C2. Consistent with above, *Prg4* expression was largely restricted to C2 among the three progenitor clusters (Fig. 1H).

At d10, the same three stem cell clusters were detected (Fig. 1F), but the relative size of C6 and C20 increased compared with d5 (Fig. 1F). Importantly, *Prg4* expression in C2 was substantially reduced at d10 relative to d5 (Fig. 1I), consistent with bulk RNA-seq data showing decreased *Prg4* expression between these timepoints (Fig. 1B). At d10, *Prg4* was also detected in soft-callus chondrocytes (C25 and C30; Supplementary Fig. 1D), but at much lower levels than in C2 (Fig. 1J). Immunofluorescence (IF) staining confirmed PRG4 protein expression in these chondrocytes (Supplementary Fig. 1E).

Together, these findings demonstrate that PSCs are the predominant source of *Prg4* during the early inflammatory phase of fracture healing, and that expression persists at later stages, albeit at reduced levels.

### The callus microenvironment induces *Prg4* expression

PDGFRα⁺SCA1⁺ cell isolation is a widely used strategy to enrich mesenchymal stem/osteochondral progenitors from callus, periosteum, or bone marrow (BM)^37,39^. Given the higher *Pdgfrα* and *Sca1* expression in C2 compared with C6 (Fig. 1E), and the fact that C2 contained ∼10-fold more cells than C6 (Fig. 1F), our data indicate that C2 represents the vast majority of PDGFRα⁺SCA1⁺ cells in the d5 callus.

To corroborate the scRNA-seq findings (Fig. 1D, H) and confirm that C2 PSCs are the primary source of *Prg4* in the d5 callus, we isolated PSCs (CD45⁻CD31⁻PDGFRα⁺SCA1⁺ cells) by fluorescence-activated cell sorting (FACS) and performed quantitative polymerase chain reaction (qPCR). *Prg4* expression was found to be largely restricted to PSCs within the healing hematoma (Fig. 2A).

**Figure 2.**
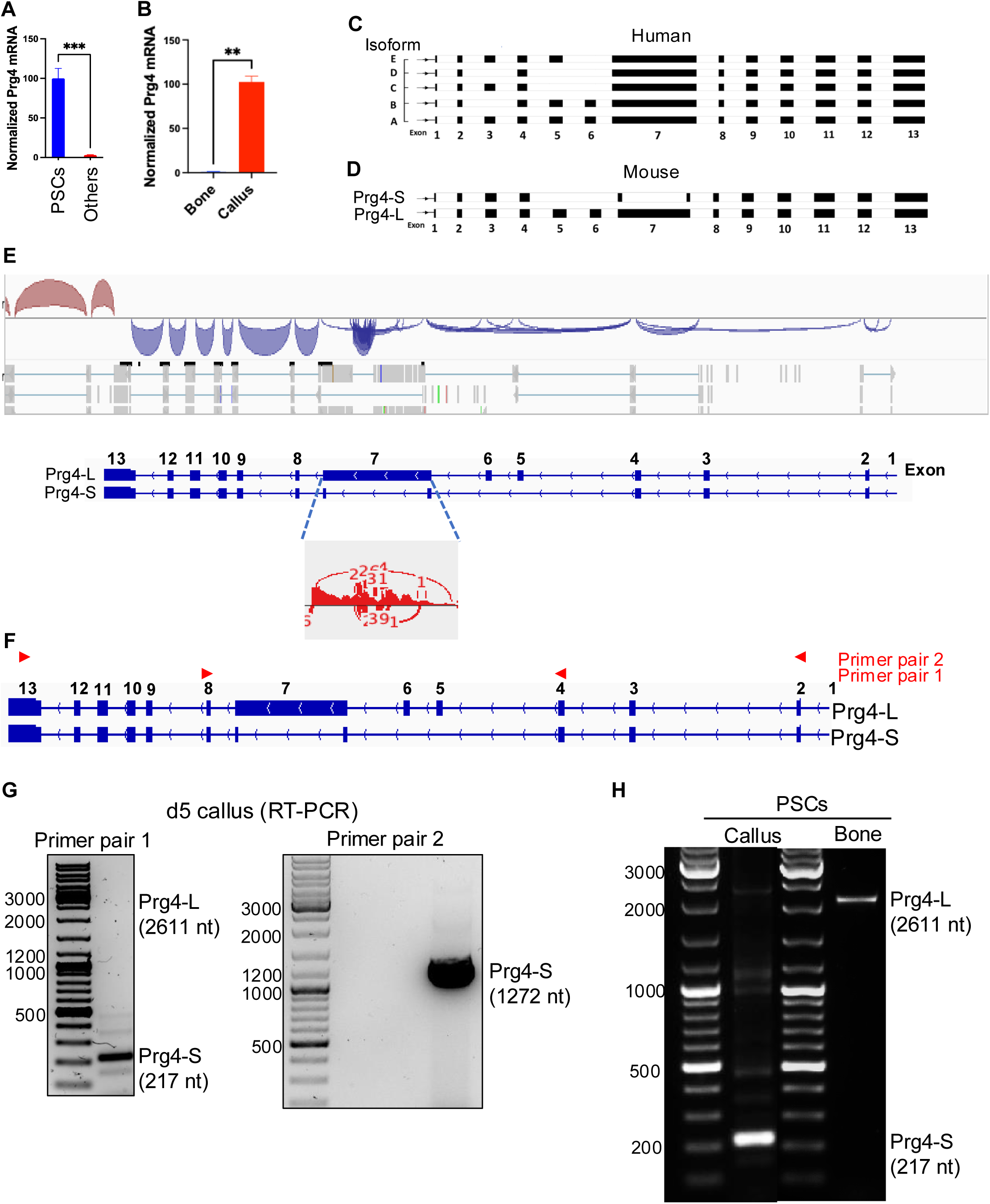
A unique isoform of *Prg4* is expressed in the callus. (**A**) RT-qPCR analysis of Prg4 mRNA in PSCs versus other cells following FACS sorting of callus cells. Prg4 mRNA levels were normalized to β-actin mRNA, with the level in PSCs defined as 100% (N = 4). (*) P < 0.001 by Welch’s test. (**B**) As in (**A**), except Prg4 mRNA levels were measured in the indicated tissues, with the level in bone defined as 1. (**\*\***) P < 0.01. (**C, D**) Exon–intron structure of human (**C**) and mouse (**D**) Prg4 pre-mRNA. Solid boxes indicate exons, and different splice isoforms are shown. (**E**) Analysis of *Prg4* splicing isoforms using bulk RNA-seq data and rMATS (top). The exon– intron structures of murine Prg4-L and Prg4-S mRNA isoforms are shown for comparison (middle). Analysis of splicing around exon 7 (gray box, bottom) shows frequent exon 7 skipping events. (**F**) Locations of Prg4 primer pairs 1 and 2. (**G**) PCR of day 5 callus RNA using Prg4 primer pair 1 (left) or 2 (right). Expected sizes of Prg4-S and Prg4-L PCR products are indicated in parentheses. (**H**) As in (**G**), except PCR was performed using RNA purified from PSCs sorted from day 5 callus or intact bone. Primer pair 1 was used. All gel images are representative of N = 3. Bar graphs show mean ± SEM.

We next asked whether the elevated *Prg4* expression was a unique property of callus PSCs or a general feature of these cells independent of their niche. Using FACS, we isolated CD45⁻CD31⁻PDGFRα⁺SCA1⁺ cells from both d5 callus and contralateral unfractured bone. qPCR revealed that *Prg4* levels were >100-fold higher in callus-derived PSCs than in their counterparts from intact bone (Fig. 2B). Consistent with these results, *Prg4* expression was also significantly enriched in total RNA from whole callus tissue compared with intact bone (data not shown).

Together, these results demonstrate that PSCs are induced to express high levels of *Prg4* in response to the inflammatory callus microenvironment during the early stages of fracture healing.

### The callus PSCs express a unique *Prg4* isoform lacking the mucin domain

The longest murine *Prg4* transcript, analogous to the longest human isoform, comprises 13 exons (Fig. 2C, D). Exon 7 encodes the central mucin-like domain, which contains numerous glycosylation sites (Fig. 2C, D; Supplementary Fig. 2). Upstream of exon 7, the transcript encodes two somatomedin-B–like domains and heparin-binding sites, whereas downstream exons encode hemopexin-like domains (Fig. 2C, D; Supplementary Fig. 2). Functionally, the heavily glycosylated mucin domain is essential for PRG4’s lubricating role at the articular cartilage surface, while the N- and C-terminal domains mediate protein–protein interactions and signaling^5–8^.

Human PRG4 (hPRG4) isoforms annotated in RefSeq, and those detected in articular cartilage^32^, lack one or more of exons 3–6 (Fig. 2C, isoforms B–E, compared to the full-length isoform A). The functions of these isoforms remain unknown, but importantly, all retain the mucin domain (Fig. 2C). In contrast, RefSeq lists two murine *Prg4* isoforms: a full-length transcript containing all exons, which we designate *Prg4*-Long (*Prg4-L*), and a shorter variant, *Prg4*-Short (*Prg4-S*), which lacks exons 5, 6, and the mucin-like repeats encoded by exon 7 (Fig. 2D). Notably, *Prg4-S* has not previously been detected or studied in musculoskeletal tissues.

To investigate isoform usage during fracture repair, we analyzed our bulk RNA-seq data^27,33^ with rMATS^40^ to identify alternative splicing events. Interestingly, *Prg4-S* was predicted to be a predominant isoform in the callus (Fig. 2E). We validated this prediction by RT-PCR of callus RNA using two primer sets spanning the skipped exons (Fig. 2F). A major band corresponding to *Prg4-S* was detected, whereas *Prg4-L* was undetectable (Fig. 2G; Supplementary Fig. 3). Sequencing confirmed the identity of the PCR product as *Prg4-S* (Supplementary Fig. 4A, B). Consistently, *Prg4-S* was the predominant isoform in callus PSCs (Fig. 2H), while, in a striking contrast, PSCs from unfractured bone primarily expressed *Prg4-L* (Fig. 2H).

Together, these results indicate that mobilization of PSCs into the callus microenvironment reprograms *Prg4* pre-mRNA processing to favor expression of *Prg4-S*, a unique isoform lacking the mucin domain whose biological function remains unexplored.

### *Prg4-S* mRNA encodes a secreted protein

The protein encoded by *Prg4-L* mRNA is known to be secreted^9–11^. We next asked whether *Prg4-S* also encodes a secreted protein. Motif analysis of PRG4-L identified the N-terminal 24 amino acids as a signal peptide directing the protein to the secretory pathway (Fig. 3A)^41,42^. Sequence alignment showed that this signal peptide is retained in PRG4-S (Fig. 3B), suggesting that PRG4-S is also secreted. To test this directly, we cultured primary PSCs sorted from the callus, collected both cells and conditioned medium after 2 days, and performed PRG4 Western blotting. PRG4-S was detected predominantly in the medium (Fig. 3C), confirming that PRG4-S is secreted.

**Figure 3.**
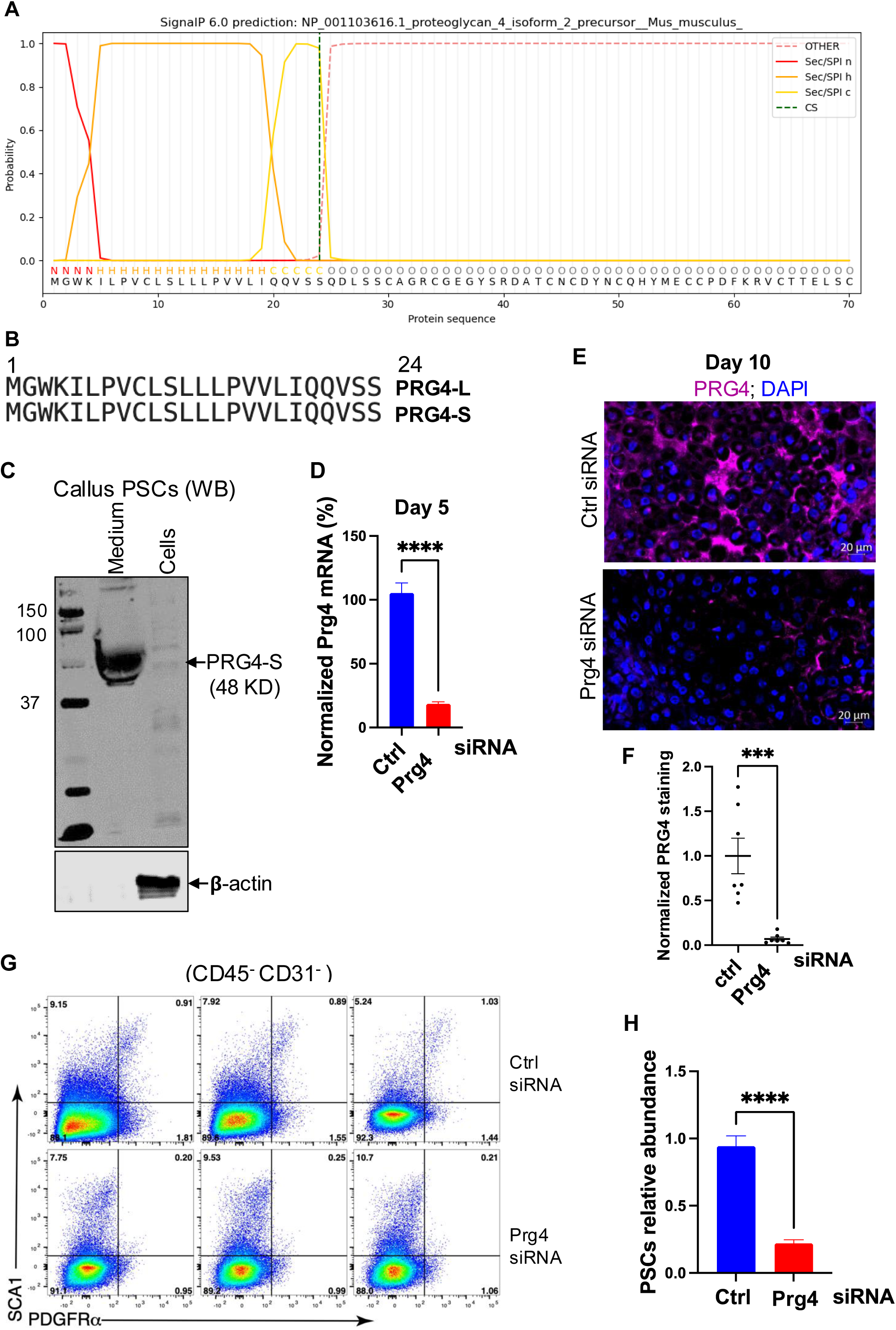
PRG4-S is a secreted protein that regulates the abundance of PSCs in the callus. (**A**) Sequence analysis of murine PRG4-L protein predicting an N-terminal signal peptide (24 amino acids). (**B**) Alignment of the N-terminal 24 amino acids in murine PRG4-L and PRG4-S proteins. (**C**) WB of PRG4 and β-actin in primary PSC cultures (isolated from day 5 callus). WB was performed on both the cells and the culture medium. The blot is representative of N = 3. (**D**) RT-qPCR of *Prg4* in RNA isolated from day 5 callus of mice treated with control (Ctrl) or Prg4 siRNA. *Prg4* mRNA levels were normalized to β-actin mRNA, with the level in Ctrl-treated mice defined as 100%. The probe used quantitated total *Prg4* expression (not isoform-specific) (N = 4). (**E**) IF staining of PRG4 (magenta) in day 10 callus of Ctrl- or Prg4 siRNA–treated mice. DAPI (blue) stains nuclei. Images were taken in the soft callus region. The PRG4 antibody recognized total PRG4 protein (not isoform-specific). (**F**) Quantification of PRG4 staining shown in (**E**). The PRG4-positive area was normalized to total callus area, with Ctrl siRNA–treated mice defined as 1 (see Materials and Methods) (N = 7). (**G**) Representative FC dot plot of PSCs (CD45^−^ CD31^−^ PDGFRα^+^ SCA1^+^ cells) in day 5 callus of Ctrl- or Prg4-S siRNA–treated mice. (**H**) Quantification of PSCs from the FC analysis shown in (**G**). PSC numbers were normalized to the total number of singlets in the callus, with Ctrl siRNA–treated mice defined as 1 (N = 3; 5 mice pooled per replicate). Bar graphs show mean ± SEM. (***) P < 0.001; (****) P < 0.0001 by Welch’s test.

### *Prg4-S* maintains PSC abundance in the healing callus

The observation that the callus microenvironment induces *Prg4-S* expression in a tissue-specific manner (Fig. 2H) prompted us to investigate its function during fracture repair. We employed an atelocollagen–nanoparticle complex to achieve sustained local delivery of *Prg4* siRNA to the callus^43–45^ (see Methods). The siRNA was designed to target *Prg4-S*. qPCR and IF confirmed efficient knockdown (KD) of *Prg4* during the first 10 days of healing (Fig. 3D–F), the window when *Prg4* expression is highest (Fig. 1B).

We next assessed the impact of *Prg4-S* KD on PSCs, its primary source. Flow cytometry (FC) at day 5 revealed an ∼5-fold reduction in PSC abundance in the KD group compared with controls (Fig. 3G, H). These findings suggest that PRG4-S promotes PSC maintenance in the callus, potentially through autocrine signaling, and thereby supports normal fracture healing.

### PRG4-S exerts paracrine immunomodulatory effects

As a secreted protein, PRG4-S may influence cellular populations within the callus through paracrine signaling. To test this, we assessed immune-cell dynamics following *Prg4-S* knockdown (KD). Consistent with our previous report of robust immune infiltration on d5^27^, *Prg4-S* KD further increased the total number of immune cells in the d5 callus by ∼30% (Fig. 4A). We next examined specific immune compartments. Among myeloid cells—the dominant immune population in the callus^27,46–54^—*Prg4-S* KD significantly increased the abundance of CD11b⁺ myeloid cells (Fig. 4B). This expansion was largely driven by polymorphonuclear granulocytes (PMNs) and myeloid progenitors, the major subsets within the CD11b⁺ myeloid population (Fig. 4C). Additional myeloid subsets were also affected: macrophages, non-classical (Ly6C^lo^) monocytes, and eosinophils were all elevated, while classical monocytes were reduced (Fig. 5A, B).

**Figure 4.**
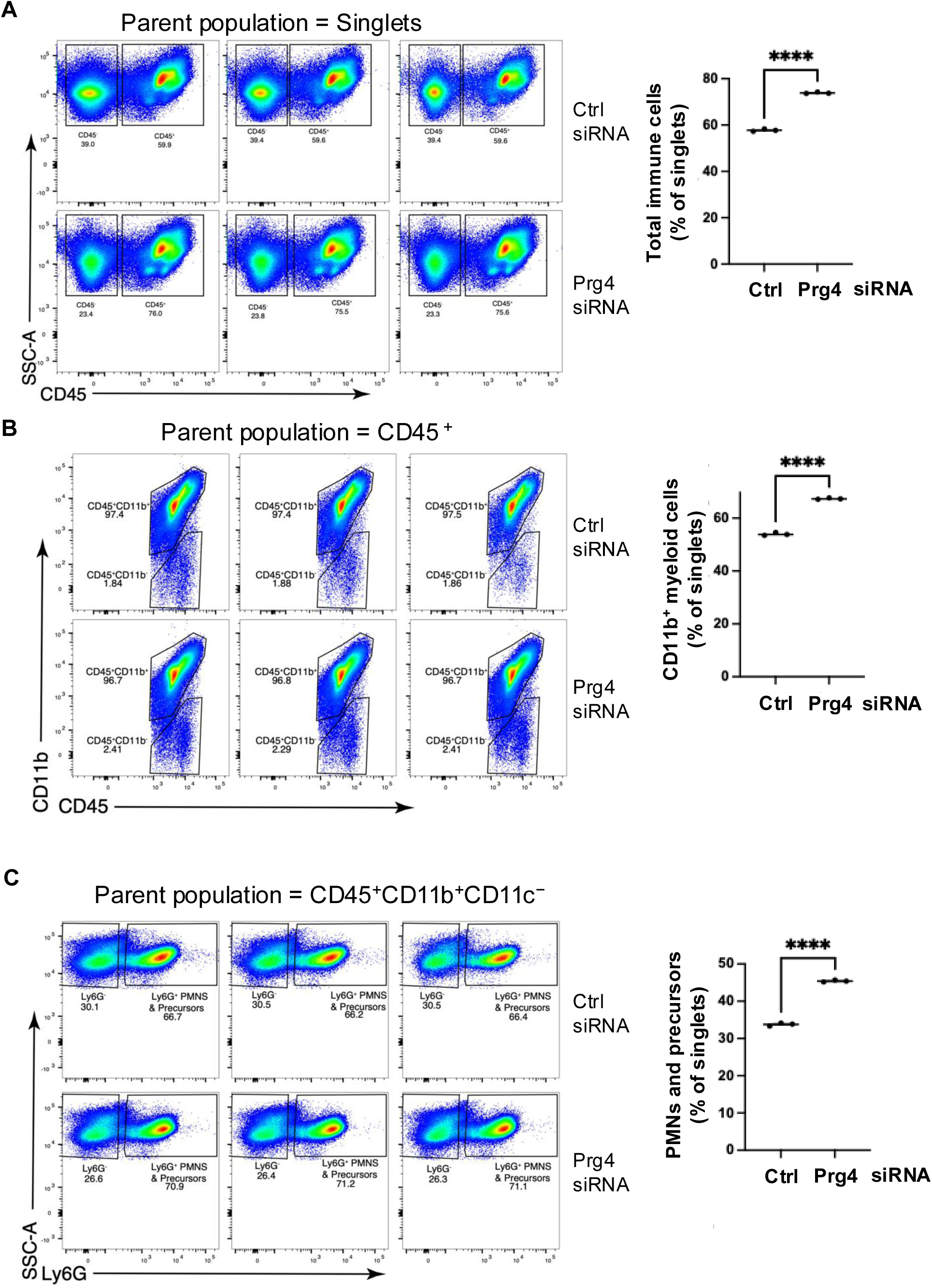
Depletion of Prg4-S increases the number of myeloid cells in the callus. (**A**) FC dot plot (left) and quantification (right) of total immune (CD45^+^) cells in day 5 callus of Ctrl- or Prg4 siRNA–treated mice. Immune cell numbers were normalized to the total number of singlets and expressed as a percentage. (**B**) As in (**A**), except myeloid (CD45^+^ CD11b^+^) cells were analyzed. (**C**) As in (**A**), except PMNs and precursors were analyzed. Notably, the numbers in the dot plots (left) represent the relative abundance of each population as a percentage of the parent population indicated in each panel, whereas the scatter plots (right) display normalized counts relative to singlets (see Supp. Table 1 for details on parent populations and gating strategies). Scatter plots show mean ± SEM. N = 3 (5 mice pooled per replicate). (****) P < 0.0001 by Welch’s test.

**Figure 5.**
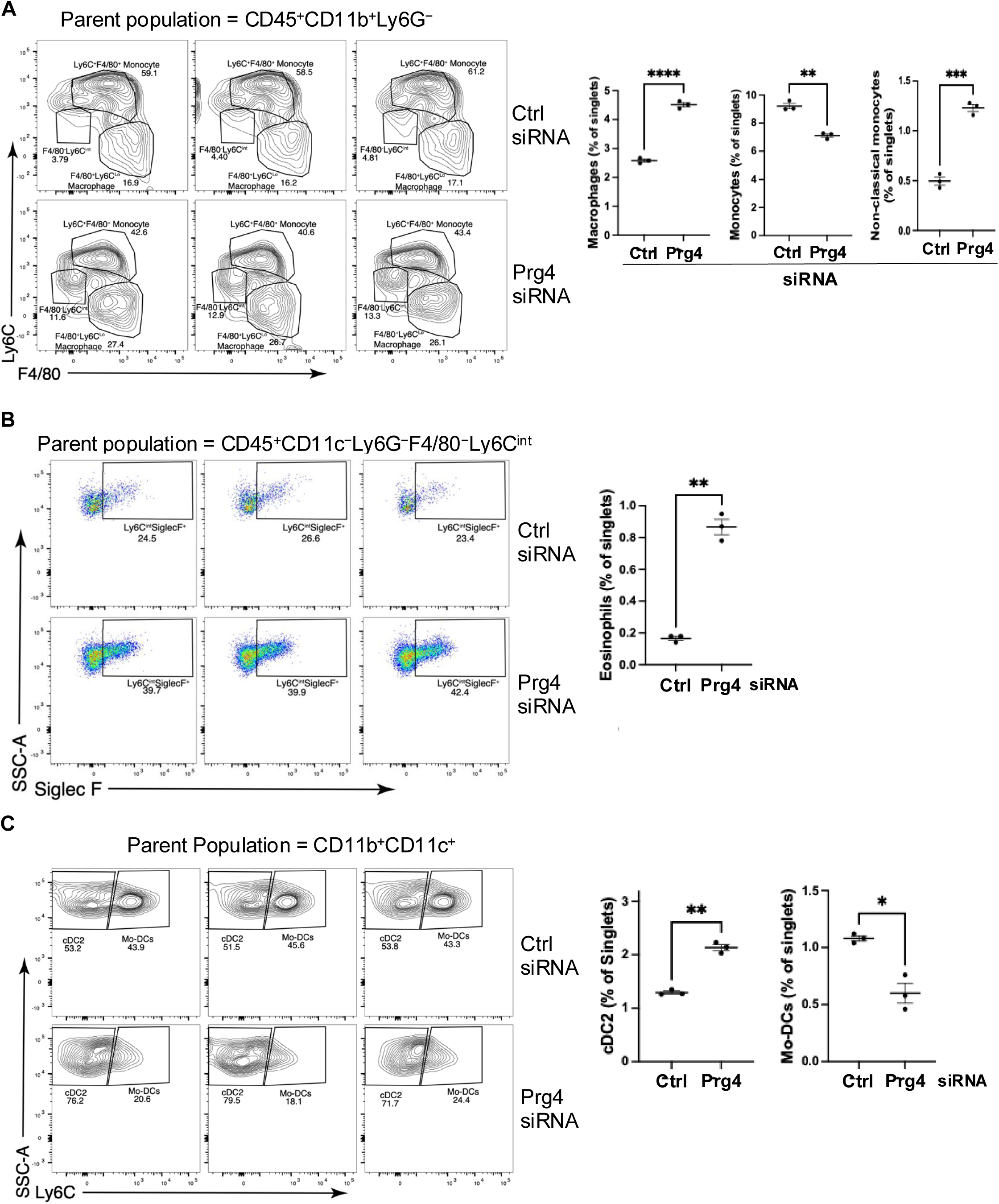
Dysregulated innate and dendritic cell responses in Prg4-S knockdown mice. (**A**) FC dot plot (left) and quantification (right) of macrophages and classical (Ly6C^Hi^) and non- classical (Ly6C^Lo^) monocytes in day 5 callus of Ctrl- or Prg4 siRNA–treated mice. Cell numbers were normalized to the total number of singlets and expressed as a percentage. (**B**) As in (**A**), except eosinophils were analyzed. (**C**) As in (**A**), except conventional DC2 (cDC2) and monocyte-derived DCs (Mo-DCs) were analyzed. Notably, the numbers in the dot plots (left) represent the relative abundance of each population as a percentage of the parent population indicated in each panel, whereas the scatter plots (right) display normalized counts relative to singlets (see Supp. Table 1 for details on parent populations and gating strategies). Scatter plots show mean ± SEM. N = 3 (5 mice pooled per replicate). (*) P < 0.05; (**) P < 0.01; (***) P < 0.001; (****) P < 0.0001 by Welch’s test.

Dendritic cells (DCs), which bridge innate and adaptive immunity^55^, displayed subset-specific responses. *Prg4-S* KD increased conventional DCs (cDC1 and cDC2), reduced monocyte-derived DCs, and left plasmacytoid DCs unchanged (Fig. 5C; Supplementary Fig. 5).

Lymphoid populations in the d5 callus^27^ were also altered by *Prg4-S* KD. In the T-cell compartment, *Prg4-S* KD selectively increased CD8⁺ T cells without affecting CD4⁺ T cells (Fig. 6A). Total B cells were elevated (Fig. 6B), with increases observed in both immature (B220^lo^) and mature (B220^hi^) subsets (Fig. 6C).

**Figure 6.**
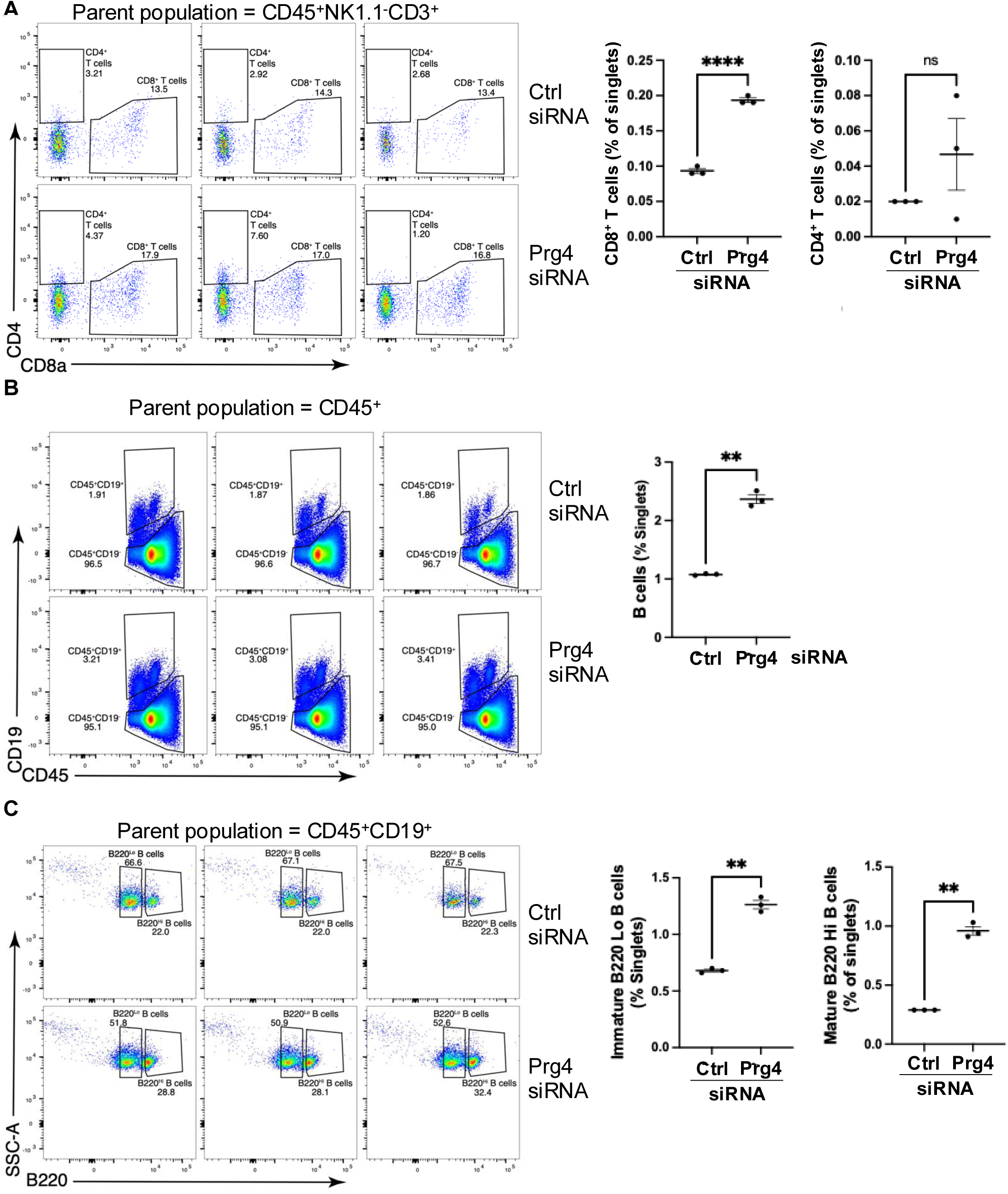
Prg4-S knockdown causes dysregulated adaptive immune responses. (**A**) FC dot plot (left) and quantification (right) of CD8^+^ and CD4+ T cells in day 5 callus of Ctrl- or Prg4 siRNA–treated mice. Cell numbers were normalized to the total number of singlets and expressed as a percentage. (**B**) As in (**A**), except total B cells were analyzed. (**C**) As in (**A**), except mature (B220^Hi^) and immature (B220^Lo^) B cells were analyzed. Notably, the numbers in the dot plots (left) represent the relative abundance of each population as a percentage of the parent population indicated in each panel, whereas the scatter plots (right) represent normalized counts relative to singlets (see Suppl. Table 1 for details on parent populations and gating strategies). Scatter plots show mean ± SEM. N = 3 (5 mice pooled per replicate). (**) P < 0.01; (****) P < 0.0001 by Welch’s test.

Together, these findings demonstrate that loss of PRG4-S exacerbates immune-cell infiltration and skews the balance of both myeloid and lymphoid compartments during the early inflammatory phase of healing.

### *Prg4-S* regulates the homeostasis of soft-callus chondrocytes

To further assess the paracrine effects of PRG4-S, we analyzed d10 callus tissue from mice treated with either control or *Prg4* siRNA. At this stage, the fracture site is bridged by a bulky cartilaginous (soft) callus containing proliferating and hypertrophic chondrocytes, surrounded by woven bone at distal sites^2,27,56^. *Prg4-S* KD caused marked defects in soft-callus formation, including reduced callus size (Fig. 7A; Supplementary Fig. 6). KD also accelerated chondrocyte hypertrophy, as indicated by increased expression of the pre-hypertrophic marker IHH (Fig. 7B) and the hypertrophic markers MMP13 and Col X (Fig. 7C, D). In addition, KD reduced chondrocyte proliferation (Fig. 7E) and increased apoptosis (Fig. 7F).

**Figure 7.**
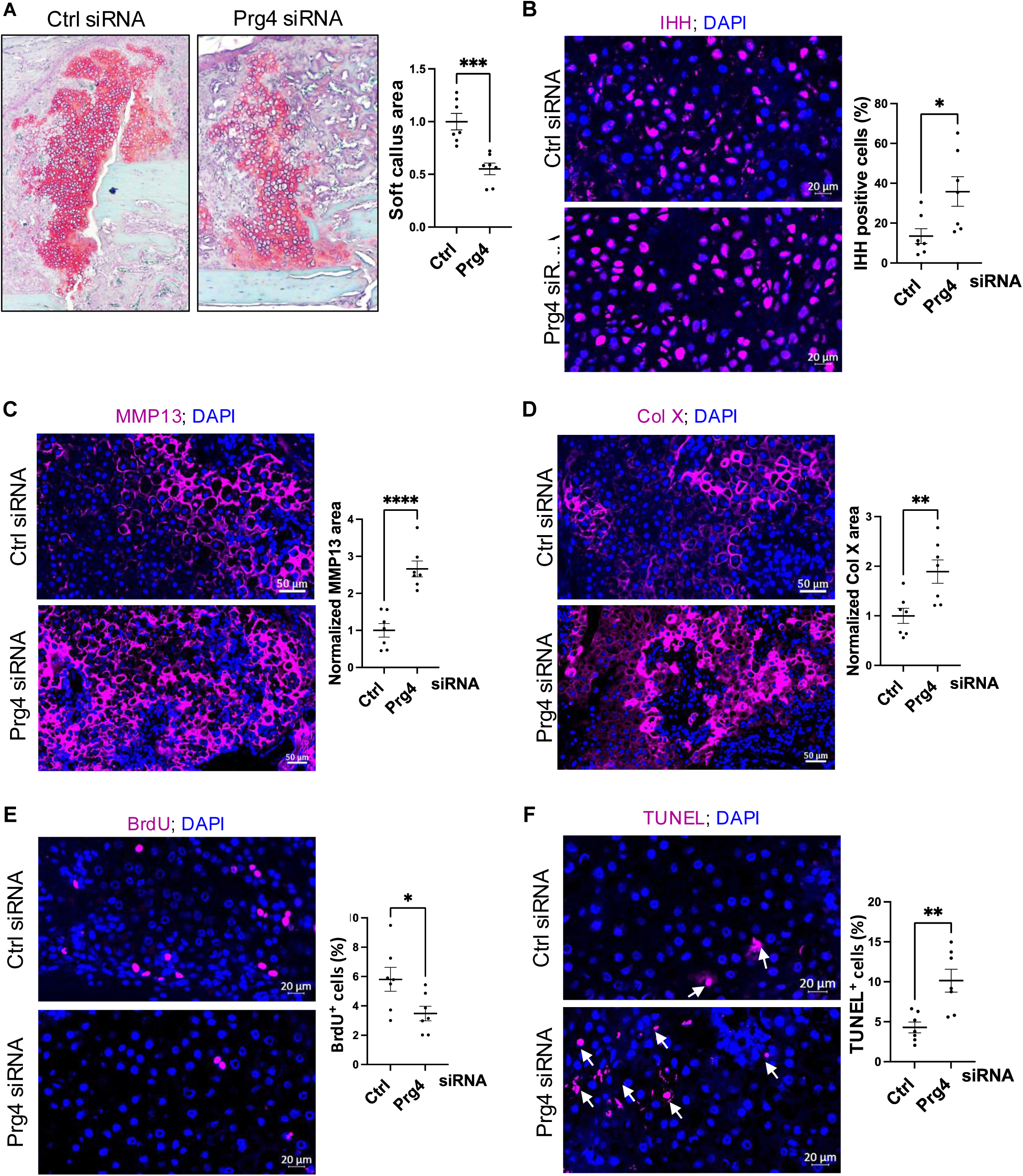
Prg4-S regulates the homeostasis of callus chondrocytes. (**A, left**) Safranin-O/fast green staining of day 10 callus from Ctrl- or Prg4 siRNA–treated mice. The soft callus (SC) is indicated in red/orange (see also **Supplementary** Fig. 6). (**A, right**) Quantification of SC area, normalized to Ctrl siRNA (set as 1). (**B, left**) IF staining of IHH (magenta) in SC; nuclei were counterstained with DAPI (blue). Scale bar, 20 μm. (**B, right**) Quantification of IHH⁺ chondrocytes as a percentage of total chondrocytes (defined by DAPI) in SC. (**C, left**) As in (**B, left**), but stained for MMP13 (magenta). Scale bar, 50 μm. (**C, right**) Quantification of MMP13⁺ area normalized to SC area, with Ctrl siRNA set as 1. (**D**) As in (**C**), but Col X was stained (left) and quantified (right). (**E**) As in (**B**), but BrdU was stained (left) and quantified (right). BrdU was injected 24 h before harvest; uptake was used as a proliferation marker. (**F, left**) As in (**B, left**), but TUNEL assay was performed to detect apoptotic cells. White arrows indicate TUNEL⁺ cells. (**F, right**) Quantification of TUNEL⁺ cells in SC as in (**B, right**). All analyses in (**B–F**) were performed in SC regions defined in (**A**) and **Supplementary** Fig. 6. Representative IF images from 7 mice are shown. Scatter plots display mean ± SEM. (*) *P* < 0.05; (**) *P* < 0.01; (***) *P* < 0.001; and (****) *P* < 0.0001 by Welch’s test.

Together, these results demonstrate that *Prg4-S* is essential for proper soft-callus formation and chondrocyte homeostasis, where it promotes proliferation while restraining premature hypertrophy and apoptosis.

### *Prg4-S* promotes new bone formation

Micro-CT (μCT) analysis of d10 callus revealed that *Prg4-S* KD significantly reduced both bone volume fraction (BV/TV) and bone mineral density (BMD) compared with control siRNA (Fig. 8A), indicating a key role for *Prg4-S* in new bone formation. Notably, *Prg4-S* KD did not alter cell proliferation or apoptosis within the woven bone compartment (Fig. 7B, C), suggesting that *Prg4-S* promotes bone formation through mechanisms beyond direct regulation of bone-cell viability.

**Figure 8.**
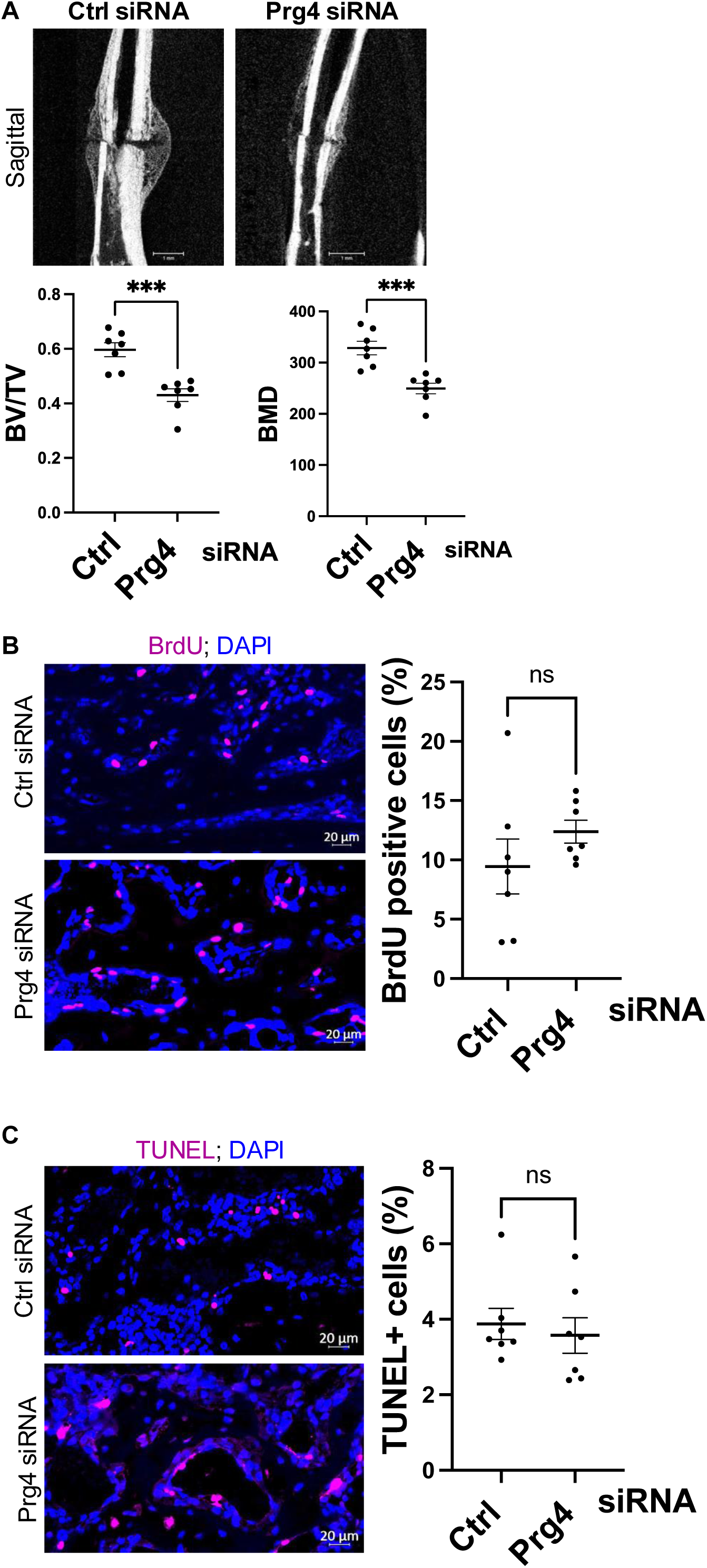
Prg4-S promotes bone formation. (**A**) μCT images (top) and quantification (bottom) of day 10 callus harvested from Ctrl- or Prg4 siRNA–treated mice. (**B, left**) IF staining of BrdU (magenta) in day 10 callus of Ctrl- or Prg4 siRNA–treated mice. BrdU was injected 24 h prior to harvest as a marker of proliferation. Staining was performed in the woven bone area (see **Supplementary** Fig. 6). DAPI (blue) stains nuclei. Scale bar, 20 μm. (**B, right**) Quantification of BrdU staining. The number of BrdU^+^ cells was normalized to the total number of cells in the woven bone area (defined by DAPI) and expressed as a percentage (see Materials and Methods). (**C, left**) As in (**B, left**), except a TUNEL assay for apoptotic cells was performed. (**C, right**) Quantification of TUNEL^+^ cells, as in (**B, right**). IF images are representative of 7 mice. Scatter plots show mean ± SEM. (***) P < 0.001 by Welch’s test.

## Discussion

The integrity of the ECM is fundamental for bone structure and mechanical strength^3^. Beyond providing structural scaffolding, ECM proteins regulate diverse cellular processes^3,57^, though many of these regulatory functions remain incompletely defined. Here, we uncover novel functions of the ECM component PRG4 and demonstrate that it orchestrates biological processes central to bone regeneration. Notably, we identify that the fracture callus microenvironment induces expression of a specific *Prg4* isoform, *Prg4-S*.

As a proteoglycan, PRG4 is highly glycosylated, with GAG chains mediating its established roles in articular cartilage, including hydration and lubrication of load-bearing surfaces^7,8^. These glycosylation sites are located in the mucin domain, encoded by exon 7 of the *Prg4-L* transcript, which comprises ∼62% of its coding sequence. In contrast, the murine *Prg4-S* isoform lacks nearly the entire mucin domain—likely explaining why it has remained unexplored. The absence of this bulky, highly glycosylated domain renders many of PRG4’s structural and lubricating functions inapplicable to PRG4-S. Instead, PRG4-S retains the N- and C-terminal regions containing protein–protein and receptor-binding motifs, which likely underlie its ability to act as a signaling molecule during callus healing. Furthermore, loss of the mucin domain may enhance diffusion, enabling PRG4-S to exert broad paracrine effects across diverse cellular populations.

Within the callus, *Prg4* is expressed by a subset of periosteal cells that display canonical markers of osteochondral progenitors, including *Postn⁺* cells^36,37^. The high relative abundance of these progenitors, particularly at early stages, aligns with prior studies demonstrating the central role of *Pdgfrα*⁺ *Sca1*⁺ and *Postn⁺* periosteal progenitors in fracture repair^36,37^. Recent reports have also identified *Prg4⁺* stem cells in other tissues that migrate to injury sites, expand, and contribute to repair^58,59^. Our results extend these findings by showing that injury induces robust *Prg4-S* expression in progenitors, where secreted PRG4-S sustains progenitors’ expansion and promotes their chondrogenic and osteogenic differentiation. This may help explain the robust regenerative potential of *Prg4⁺*stem cell populations. More broadly, *Prg4* exemplifies a class of injury-inducible genes that are minimally expressed during homeostasis but strongly upregulated following tissue damage.

The induction of *Prg4-S* during callus formation reflects an isoform switch that favors PRG4-S over PRG4-L. This guided our efforts to define isoform-specific roles during healing. Previous studies showed that PRG4-L reduces macrophage recruitment, forming the basis for its therapeutic use in osteoarthritis^22,23,60^. Here, we show that PRG4-S also regulates immunity, but in the fracture context: loss of PRG4-S led to exaggerated immune infiltration, primarily due to expansion of myeloid populations, along with an increased CD8⁺/CD4⁺ T-cell ratio—both features of inflammatory states associated with impaired healing^27,61^. These findings reveal that PRG4-S exerts immunomodulatory functions that might be distinct from those of PRG4-L and suggest that dysregulation of these effects contributes to defective repair in *Prg4-S*–depleted mice. Some of these immune effects may be direct, whereas others likely arise through intercellular cross-talk. Future studies should dissect isoform-specific immune pathways and determine the extent of functional overlap between PRG4-S and PRG4-L.

Beyond immunity, PRG4-S shares chondrogenic and chondroprotective properties with PRG4-L, such as inhibition of chondrocyte hypertrophy. However, their physiological contexts diverge: PRG4-L acts in articular cartilage, where hypertrophy must be suppressed to prevent osteoarthritis, while PRG4-S functions in fracture callus, where controlled hypertrophy drives endochondral ossification. Because PRG4-L is not expressed in the callus, fracture healing does not permit direct comparison of isoform-specific contributions. Studies in tissues where both isoforms are co-expressed will be essential to evaluate whether individual isoforms—or combinations—offer maximal therapeutic benefit.

The receptors mediating PRG4-S signaling remain undefined. Prior work suggested interactions between full-length PRG4 and Toll-like receptors or CD44 in the synovium^62–64^, but these observations are limited. Given its structural differences, PRG4-S may engage distinct receptors. Considering the cellular heterogeneity of the callus and the wide-ranging effects of PRG4-S, receptor interactions are likely cell type–specific. Identifying PRG4-S receptors will be critical for designing receptor-targeted modulators to fine-tune cellular responses and optimize healing outcomes.

In summary, we establish PRG4-S as a novel pro-regenerative isoform that is essential for fracture healing. This work shifts the focus from previously described isoforms, which are downregulated during repair, to PRG4-S as a key regulator of progenitor expansion, lineage differentiation, and immune homeostasis. Its shorter transcript, smaller protein size, and reduced structural complexity relative to mucin-containing isoforms facilitate its production and therapeutic application as either an RNA- or protein-based therapy. Collectively, our findings position PRG4-S as a promising candidate for clinical translation in the treatment of impaired fracture healing and nonunion.

## Acknowledgments

This work was supported by the National Institutes of Health (NIH) grants R01 DK121327 to R.A.E. The services and instruments of the Genome Sciences Core (RRID:SCR_021123) used in this project were funded in part by the Pennsylvania State University College of Medicine, through the Office of the Vice Dean for Research and Graduate Studies, and by the Pennsylvania Department of Health using Tobacco Settlement Funds (CURE). FC analysis and FACS were performed at the Flow Cytometry and Cell Sorting Core at the Penn State University College of Medicine (RRID:SCR_021134).

## Contributions

Study design: RAE. Data collection: MT, DKK, and RAE. Data analysis and interpretation: MT, DKK, CCN, FK, and RAE. Manuscript writing: MT, FK, and RAE.

