## Supplementary material for "A Novel Proteoglycan-4 Isoform Drives Skeletal Regeneration": Supp Figures

Supp. Fig. 1

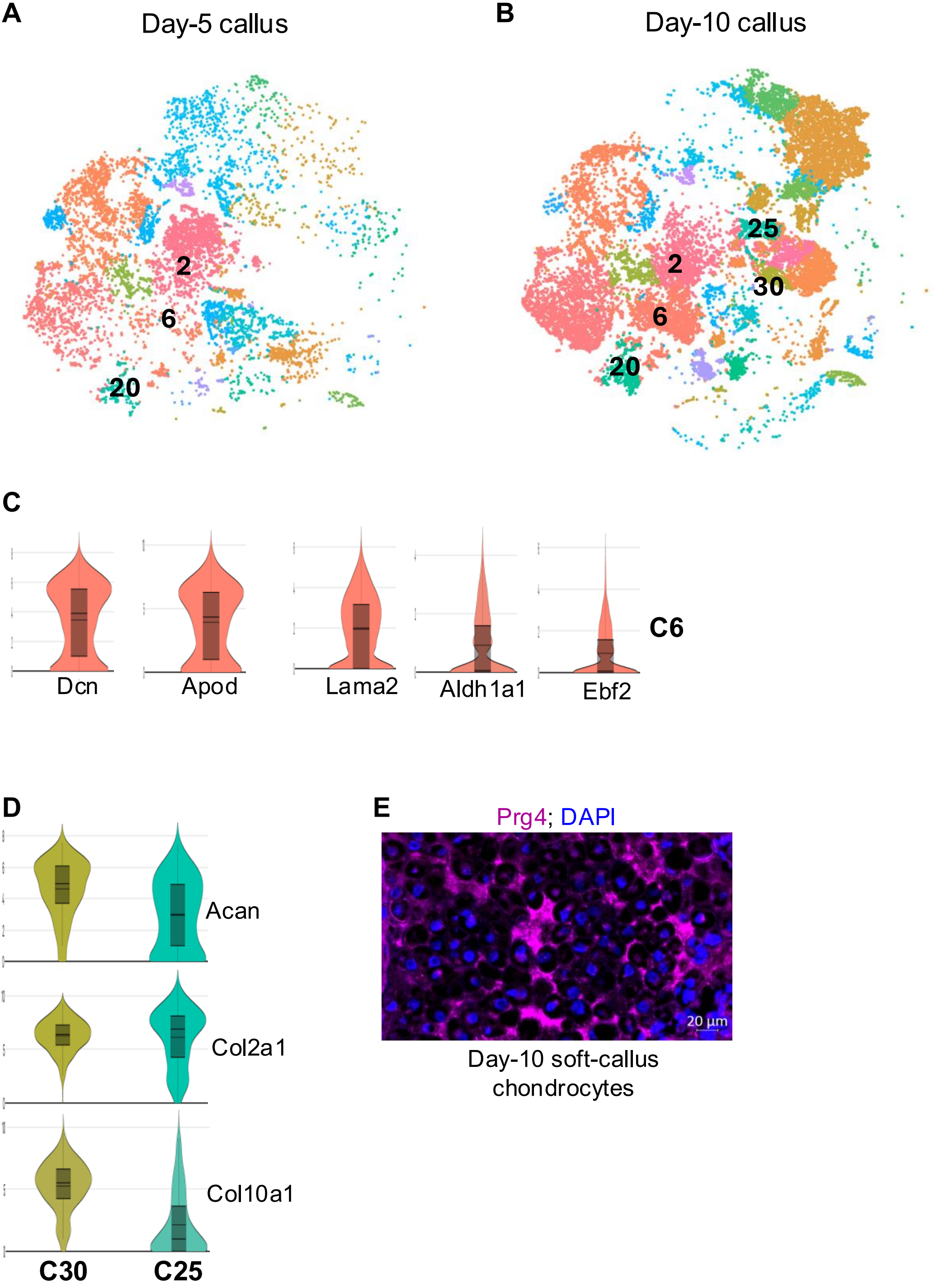
A
Day-5 callus
2
6
20
B
Day-10 callus
25
2
30
6
20
C
Dcn
Apod
Lama2
Aldh1a1
Ebf2
C6
D
Acan
Col2a1
Col10a1
C30
C25
E
Prg4; DAPI
20 µm
Day-10 soft-callus chondrocytes

**Supp. Fig. 2**

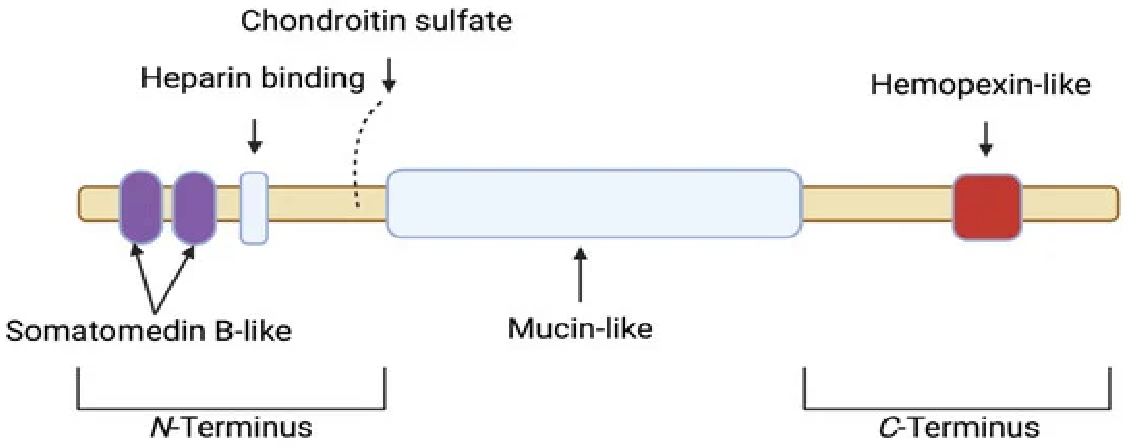

Chondroitin sulfate
Heparin binding
Hemopexin-like
Somatomedin B-like
Mucin-like
*N*-Terminus
*C*-Terminus

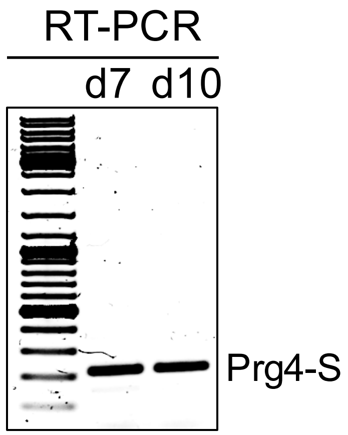
RT-PCR
d7
d10
Prg4-S

**A** 

NNNNNNNNNNNNGGCGATTGGGCCCTCTAGATGCATGCTCGAGCGGCCGCCAGTGTGATGGATAT
CTGCAGAATTCGCCCTTGTGCAGATTATGACAGCTTCTGTGAAGAAGTAAAAGATAACAAGAAAAACA
CTCCTAAAAAGAAACCCAATCCAGAA
CCGCCGGCTGTGGATGAAGCTGGAAGCGGGCTGGATAATGCTATTCCAGGCACTGATCTTTTGGCC
GGGAGACTCAATCA
AGGCATTAACATCAATCCCATGCTTTCAGATGAGACCAATTTATGCAATGGTAAGCCAGTAAGGGCGA
ATTCCAGCACAC
TGGCGGCCGTTACTAGTGGATCCGAGCTCGGTACCAAGCTTGATGCATAGCTTGAGTATTCTATAGT
GTCACCTAAATAG
CTTGGCGTAATCATGGTCATAGCTGTTTCCTGTGTGAAATTGTTATCCGCTCACAATTCCACACAACA
TACGAGCCGGAA
GCATAAAGTGTAAAGCCTGGGGTGCCTAATGAGTGAGCTAACTCACATTAATTGCGTTGCGCTCACT
GCCCGCTTTCCAG
TCGGGAAACCTGTCGTGCCAGCTGCATTAATGAATCGGCCAACGCGCGGGGAGAGGCGGTTTGCGT
ATTGGGCGCTCTTC
CGCTTCCTCGCTCACTGACTCGCTGCGCTCGGTCGTTCGGCTGCGGCGAGCGGTATCAGCTCACTC
AAAGGCGGTAATAC
GGTTATCCACAGAATCAGGGGATAACGCANGAAAGAACATGTGAGCAAAAGGCCAGCAAAAGGCCA
GGAACCGTAAAAAGGCCGCGTTGCTGGCGTTTTTCCATAGGCTCCGCCCCCCTGACGAGCATCACA
AAAATCGACGCTCAAGTCAGAGGTGGCGAAACCCGACAGGACTATAAAGATACCAGGCGTTTCCCCC
TGGNAGCTCCCNCGTGCGCTCTCCTGTTCCNACCCTGCCGC
TTACCGGATACCTGTCCGCCTTTCTCCCTNCGGGAAGCGNNGNGCTTTCTCATAGCTCACGCTGTAG
GTATCNNCANNN

**B**

```
Query: Mus musculus proteoglycan 4 (megakaryocyte stimulating factor, articular superficial zone
protein) (Prg4), transcript variant 2, mRNA Query ID: NM_001110146.1 Length: 1884


>
Sequence ID: Query_55587 Length: 1039
Range 1: 83 to 300

Score:394 bits(436), Expect:4e-113,
Identities:218/218(100%),  Gaps:0/218(0%), Strand: Plus/Plus

Query  329   TGTGCAGATTATGACAGCTTCTGTGAAGAAGTAAAAGATAACAAGAAAAACACTCCTAAA  388
             ||||||||||||||||||||||||||||||||||||||||||||||||||||||||||||
Sbjct  83    TGTGCAGATTATGACAGCTTCTGTGAAGAAGTAAAAGATAACAAGAAAAACACTCCTAAA  142

Query  389   AAGAAACCCAATCCAGAACCGCCGGCTGTGGATGAAGCTGGAAGCGGGCTGGATAATGCT  448
             ||||||||||||||||||||||||||||||||||||||||||||||||||||||||||||
Sbjct  143   AAGAAACCCAATCCAGAACCGCCGGCTGTGGATGAAGCTGGAAGCGGGCTGGATAATGCT  202

Query  449   ATTCCAGGCACTGATCTTTTGGCCGGGAGACTCAATCAAGGCATTAACATCAATCCCATG  508
             ||||||||||||||||||||||||||||||||||||||||||||||||||||||||||||
Sbjct  203   ATTCCAGGCACTGATCTTTTGGCCGGGAGACTCAATCAAGGCATTAACATCAATCCCATG  262

Query  509   CTTTCAGATGAGACCAATTTATGCAATGGTAAGCCAGT  546
             ||||||||||||||||||||||||||||||||||||||
Sbjct  263   CTTTCAGATGAGACCAATTTATGCAATGGTAAGCCAGT  300
```

**Supp. Fig. 5**

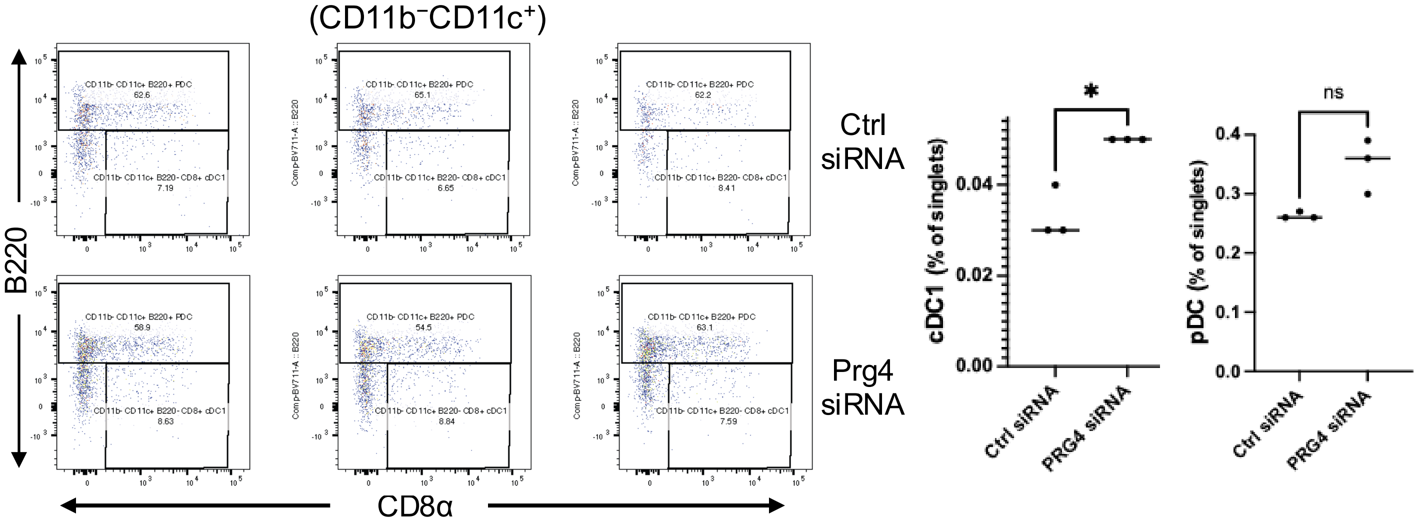
(CD11b⁻CD11c⁺)
B220
CD8α
Ctrl siRNA
Prg4 siRNA
CD11b- CD11c+ B220+ PDC
62.6
CD11b- CD11c+ B220- CD8+ cDC1
7.19
65.1
6.65
62.2
8.41
58.9
8.63
54.5
8.84
63.1
7.59
Comp-BV711-A :: B220
cDC1 (% of singlets)
0.00
0.02
0.04
*
pDC (% of singlets)
0.0
0.1
0.2
0.3
0.4
ns
Ctrl siRNA
PRG4 siRNA

A

Supp. Fig. 6

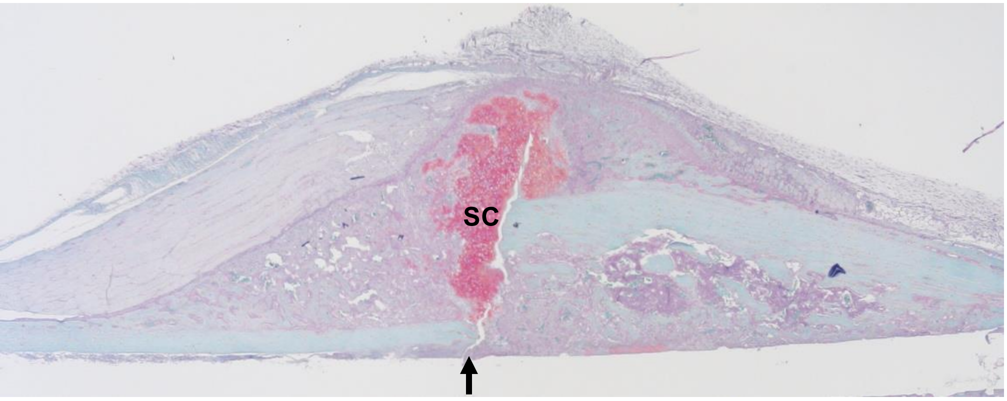


Ctrl siRNA

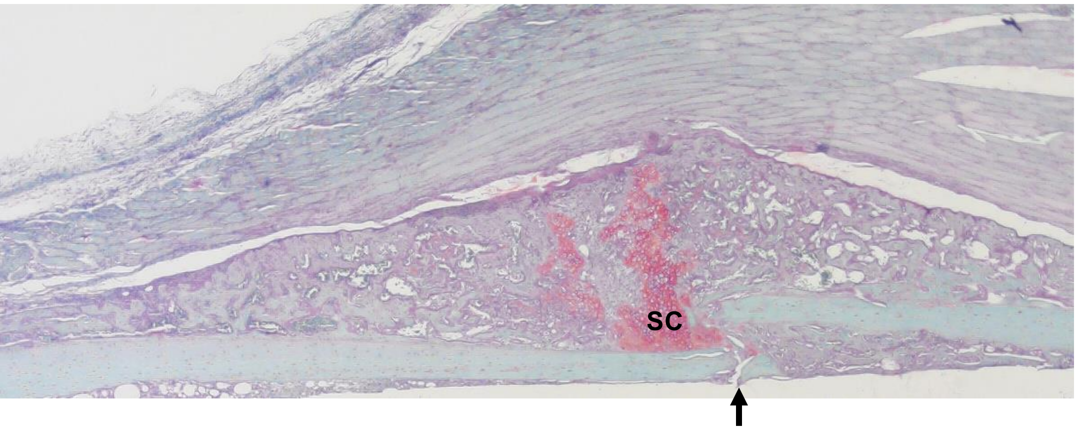


Prg4 siRNA
